# BXD51: A Robust and Translational Mouse Model for Studying the Pathophysiology of Glaucoma

**DOI:** 10.64898/2026.07.21.739658

**Authors:** Linan Guan, Xiangdi Wang, Raven Simpson, Shruthivani Velrajan, Benton Chuter, Lu Lu, Robert W. Williams, William White, T.J. Hollingsworth, Monica M. Jablonski

## Abstract

The aim of this study was to characterize the BXD51 mouse strain as a reproducible model of chronic progressive glaucoma. Unlike the highly susceptible DBA/2J (D2) mutant strain, BXD51 is a genetically stable recombinant inbred line derived from C57BL/6J (B6) and D2 parental lines. Longitudinal assessments of intraocular pressure (IOP), visual acuity (VA), contrast sensitivity (CS), and pattern electroretinogram (pERG) demonstrated that BXD51 mice undergo a delayed decline in visual and retinal ganglion cell (RGC) function. Their decline is biphasic, with a period of initial ocular stress followed by a late-onset, accelerated structural and functional deterioration of RGCs. Anterior segment structural analysis by optical coherence tomography (OCT) and histology demonstrated increasing pigment dispersion and subsequent iridocorneal angle closure. Immunofluorescence analysis of structural neuronal markers (TUBB3 and MAP1A/2) exhibited thinning of the ganglion cell layer (GCL) and inner plexiform layer (IPL) together with axonal degeneration, mirroring the laminar degeneration seen in human glaucoma patients. BXD51 also revealed marked spatial heterogeneity between peripheral and central retina. Multivariate analysis confirmed that BXD51 follows a distinct clinical trajectory that separates it from both wild-type (B6) and a severe glaucoma model (D2). By spanning the range between resistance and extreme susceptibility to glaucomatous neurodegeneration, this study establishes the BXD51 mouse as a translational platform for mechanistic studies and for evaluating long-term neuroprotective strategies.

## Introduction

Glaucoma, a vision disorder characterized by optic neuropathy, is the leading cause of irreversible blindness worldwide [1, 2]. Despite its prevalence, the molecular mechanisms driving glaucoma pathophysiology remain incompletely defined. Research has long been limited by a shortage of translationally relevant animal models, and to date no single model fully recapitulates the diverse clinical and pathologic features of human glaucoma. Inherited models of glaucoma also tend to show greater phenotypic heterogeneity in onset and progression than induced models [3, 4].

The DBA/2J (D2) mouse is a classical model of pigmentary glaucoma, in which mutations in *Gpnmb* and *Tyrp1* lead to iris pigment dispersion and subsequent elevation of intraocular pressure (IOP) and degeneration of axons in the optic nerve (ON). Although D2 mice provide a useful genetic framework, glaucoma in this strain is phenotypically variable, therefore large sample sizes are needed to reach statistical power. This variability makes it difficult to dissect the contributions of individual core and modifier genes, which limits the utility of the D2 mouse as a direct translational platform for human glaucoma even though it remains valuable as a model of pigmentary dispersion [5–8].

To address these limitations, we turned to the BXD recombinant inbred family of mice, derived from C57BL/6J (B6) and D2 parental strains [9]. An advantage of a genetically diverse family over non-inbred strains is that each strain is homozygous across its genome and can provide an essentially unlimited source of genetically identical test subjects. The breeding scheme used to generate the BXD genetic reference panel—derived from reciprocal crosses between the glaucoma-susceptible DBA/2J (D2) and the glaucoma-resistant C57BL/6J (B6) strains—allows the D2-derived, glaucoma-associated alleles to resegregate against the stable, reproducible genetic background contributed by B6, filtering out much of the variability of the parental D2 strain while retaining some of the underlying pathogenic alleles [10–13]. In earlier large-scale phenotyping work, our laboratory evaluated ≥65 BXD strains for quantitative ocular features, including IOP, ON axon counts, and iris transillumination scoring across five age cohorts [14–20], and identified BXD51 as a promising spontaneous model of glaucoma.

Building on these data, we present a multidimensional characterization of the BXD51 model. By combining longitudinal IOP, visual acuity (VA), contrast sensitivity (CS), and pattern electroretinogram (pERG) assessments with evaluation of iridocorneal angle status, *in vivo* imaging, and histology, we mapped the onset and progression of the glaucomatous hallmarks of this model. BXD51 mice present with a biphasic phenotype comprised of a well-defined latent state followed by a predictable decline at later ages. In the D2 model, by contrast, these stages are often compressed or occur unpredictably. Our data establish BXD51 as a translationally viable preclinical platform. The strain offers an extended clinical window, making it a useful tool for studying the early stages of glaucoma before permanent vision loss occurs.

## Results

Historical phenotypic data for the BXD51 strain are summarized in Supplementary Fig. 1. These records present a parallel progression of age-related IOP elevation and increased necrotic ON axon counts, with no iris transillumination defects until 13 months of age. This foundational evidence describes the ocular health trajectory that informs the current study, which extends these findings through longitudinal assessments of functional and morphological markers, serving as the primary basis for our further investigation of the BXD51.

### Longitudinal Characterization of Intraocular Pressure (IOP) and Ocular Function

Based on the outcomes of our historic data, we carried out a longitudinal assessment of the BXD51 strain to define the trajectory of its ocular phenotype. To capture the temporal evolution of glaucoma hallmarks, we monitored IOP across five cohorts ranging from 1 to 12 months of age, with B6 and D2 parental strains as comparators. B6 mice maintained a stable baseline of approximately 15-18 mmHg throughout the study period (Fig. 1A). BXD51 mice, by contrast, held a stable IOP baseline during the first 6 months of life, followed by a non-linear rise toward higher IOP levels beginning at 9 months of age. By 12 months, peak individual IOP values were comparable to, and often exceeded, the peak values recorded in the D2 strain. Although BXD51 mice varied in the timing and magnitude of pressure elevation, the 9-to-12-month interval consistently captured the hypertensive phase (Fig. 1B). The D2 strain presented with an IOP elevation that began earlier, with a broad distribution reflecting its known phenotypic heterogeneity (Fig. 1C). Together, these patterns provide a defined framework for investigating pressure-dependent pathological changes across these models.

**Figure 1.**
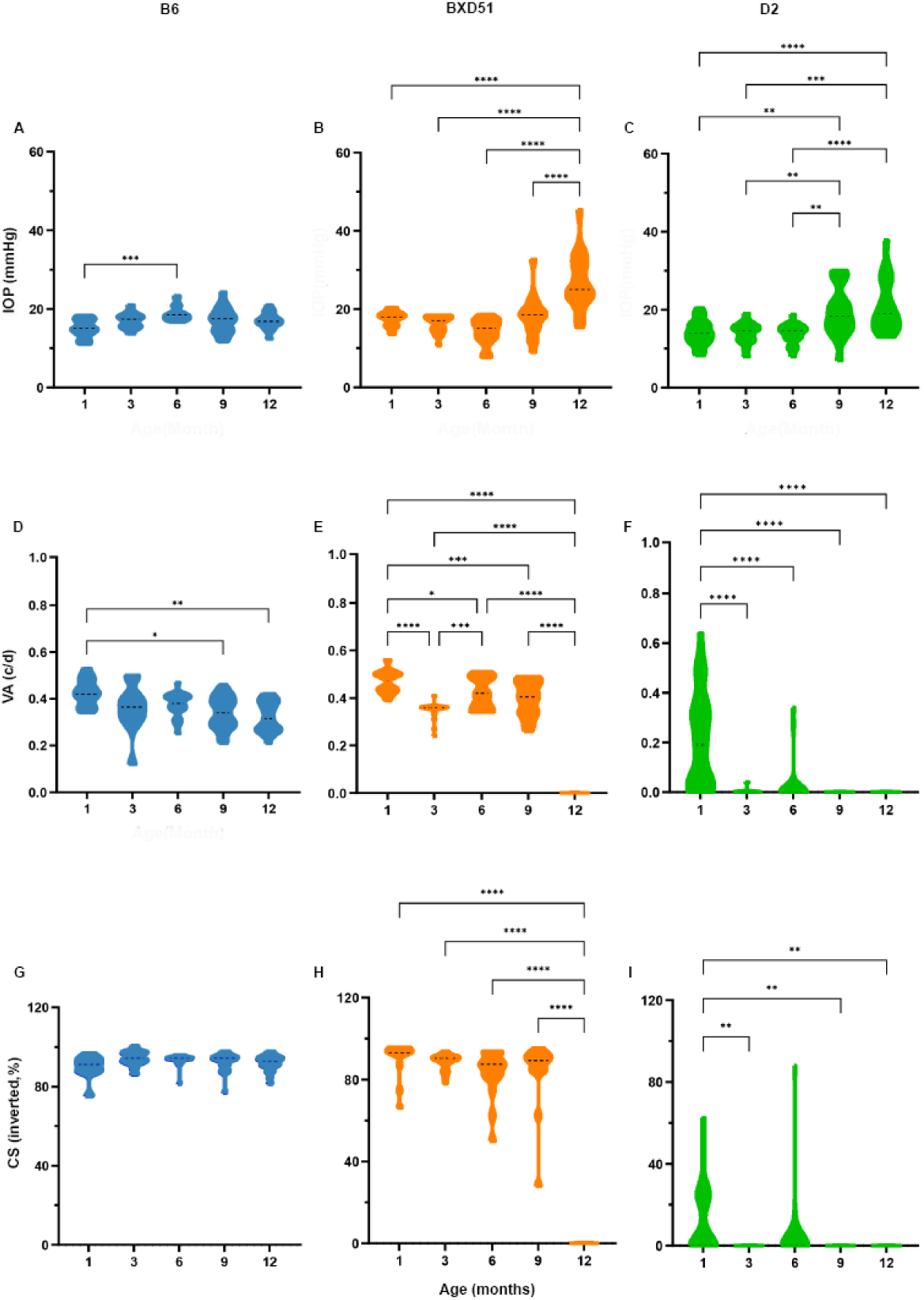
Longitudinal assessment of intraocular pressure (IOP) and visual function in B6, BXD51, and D2 mice. (A–C) Violin plots presenting IOP measurements in B6 (A), BXD51 (B), and D2 (C) mice from 1 to 12 months of age. B6 mice exhibit stable IOP throughout the observed period, whereas significant IOP elevation is observed in BXD51 and D2 mice beginning at 12 months and 9 months, respectively. (D–F) Longitudinal evaluation of visual acuity (VA), measured in cycles per degree (c/d). B6 mice (D) maintain a stable functional plateau with no significant decline until age-related senescence at 9–12 months. BXD51 mice (E) demonstrate a transient fluctuation in VA at 3 months, followed by a compensatory period of functional stability between 6 and 9 months, and ultimately a complete loss of measurable VA by 12 months. D2 mice (F) display a near-total drop in VA as early as 3 months that remains unaltered up to 12 months. (G–I) Contrast sensitivity (CS) is presented as inverted percentages. B6 mice (G) show stable CS across all time points, BXD51 mice (H) exhibit a later but significant decline by 12 months, and D2 mice (I) show early visual impairment. Data are presented as individual values within violin plots. \**P* < 0.05, \*\**P* < 0.01, \*\*\**P* < 0.001, \*\*\*\**P* < 0.0001.

We then evaluated the effect of this hypertensive profile on VA and CS at multiple time points, allowing us to relate the progression of ocular hypertension to changes in visual function. Regarding VA, B6 showed a subtle, age-dependent decline (Fig. 1D), whereas the BXD51 and D2 strains followed distinct, non-overlapping dysfunction kinetics (Figs. 1E, 1F). Progression in BXD51 mice was protracted; after a transient reduction in VA at 3 months, VA recovered and stabilized between 6 and 9 months, maintaining a compensatory plateau before a sharp late stage decline at 12 months (*P* < 0.0001, Fig. 1E). This decline preceded the onset of IOP increase, suggesting that the early pathogenesis is not simply mechanical injury secondary to high IOP but instead involves an IOP-independent upstream mechanism. In D2 mice, significant functional impairment appeared much earlier, already present by 3 months (Fig. 1F).

B6 controls maintained CS with no discernible change over time (Fig. 1G). Unlike the early changes seen in VA, CS in BXD51 remained at a plateau through 9 months and then deteriorated rapidly (*P* < 0.0001, Fig. 1H). In D2 mice, the degeneration of CS and VA was closely synchronized, dropping to near-zero levels as early as 3 months of age, with only negligible, transient traces detected thereafter (Fig. 1I).

Together, these assessments demonstrate that BXD51 behaves as a biphasic glaucoma model. The compensatory fluctuation in VA begins before any measurable increase in IOP, indicating an early, non-mechanical phase of retinal stress. The uncoupling of CS and VA at 3 months also raises questions about the underlying pathogenic mechanisms.

### Retina Ganglion Cell (RGC) Function Profiles Across Different Stages of Chronic Intraocular Hypertension

To assess RGC functional integrity, we recorded pERG responses across all three strains. B6 controls showed minor, age-related fluctuations in both P50 and N95 over the 12-month period (Figs. 2A, D). During early life (1-6 months), the RGC functional output of BXD51 mice not only remained intact but exceeded that of age-matched B6 mice. As IOP rose sharply between 9 and 12 months, however, the BXD51 cohort deteriorated. By the 12-month advanced stage of disease, both P50 (*P* < 0.0001) and N95 (*P* < 0.001) amplitudes were sharply reduced. BXD51 individuals showed wide variability at late stages (9-12 months), paralleling the broad distribution of their extreme IOP values (Figs. 2B, E). The D2 strain, in contrast, had a lower functional baseline from the outset and showed a steady worsening of RGC function. Compared with baseline at 1 month, both P50 and N95 amplitudes declined significantly; this trend emerged at 6 months and became highly significant by 9 months (*P* < 0.0001, Figs. 2C, 2F). By 12 months, the RGC response in D2 mice was almost completely extinguished, consistent with the rapid visual loss seen in the behavioral assays.

**Figure 2.**
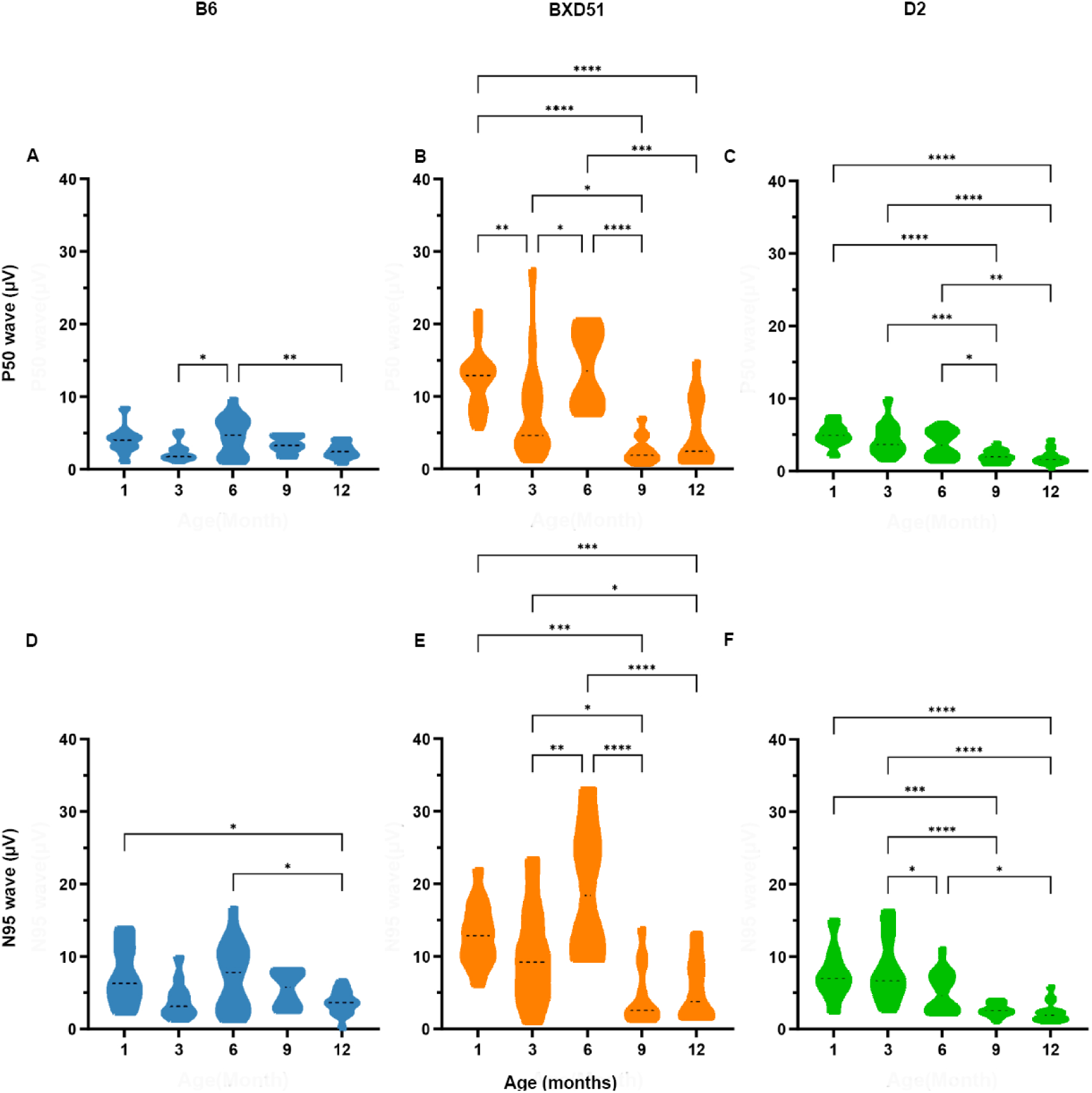
Longitudinal assessment of retinal ganglion cell function via pattern electroretinogram (pERG). (A–C) Violin plots representing the amplitude of the pERG P50 wave (µV) in B6 (A), BXD51 (B), and D2 (C) mice from 1 to 12 months of age. (D–F) Quantitative evaluation of the pERG N95 wave amplitude (µV) in the respective strains. B6 mice maintain stable pERG responses throughout the study duration, whereas BXD51 mice are characterized by a transient compensatory surge in amplitude before 6 months that surpasses B6 levels, followed by a rapid decline. In contrast, D2 mice show a steady, age-related attenuation in pERG amplitude. Data are presented as individual values within violin plots. \*\*\**P* < 0.001, \*\*\*\**P* < 0.0001.

This difference in early-stage kinetics between the two glaucoma models suggests that although RGC functional integrity and behavioral vision may follow distinct compensatory timelines, both point to a strain-dependent capacity to temporarily buffer the degenerative process.

### Characterization of Anterior Chamber Angle

To evaluate the morphological changes in the anterior chamber, we investigated all three mouse strains (B6, BXD51, D2) using OCT imaging, histology (H&E staining), and longitudinal quantification of iridocorneal angle closure.

As expected, the iridocorneal angles in the B6 group remained open, with clear separation between the iris and the posterior corneal surface (Figs. 3A, D, G). Both BXD51 and D2 mice, by contrast, showed varying degrees of angle closure. At 12 months of age, BXD51 mice showed adhesion between the iris and cornea (Figs. 3B, E, H), but these anterior synechiae were less severe than those in age-matched D2 mice (Figs. 3C, F, I). Compared with the D2 strain, BXD51 mice also showed less iris pigment dispersion, sparse pigmented deposits on the corneal endothelial surface, iris thinning and atrophy, a pupillary exudative membrane, and only mild pupil constriction. Clinically, BXD51 showed a clear lag relative to D2 in progressive angle narrowing and eventual synechial closure.

**Figure 3.**
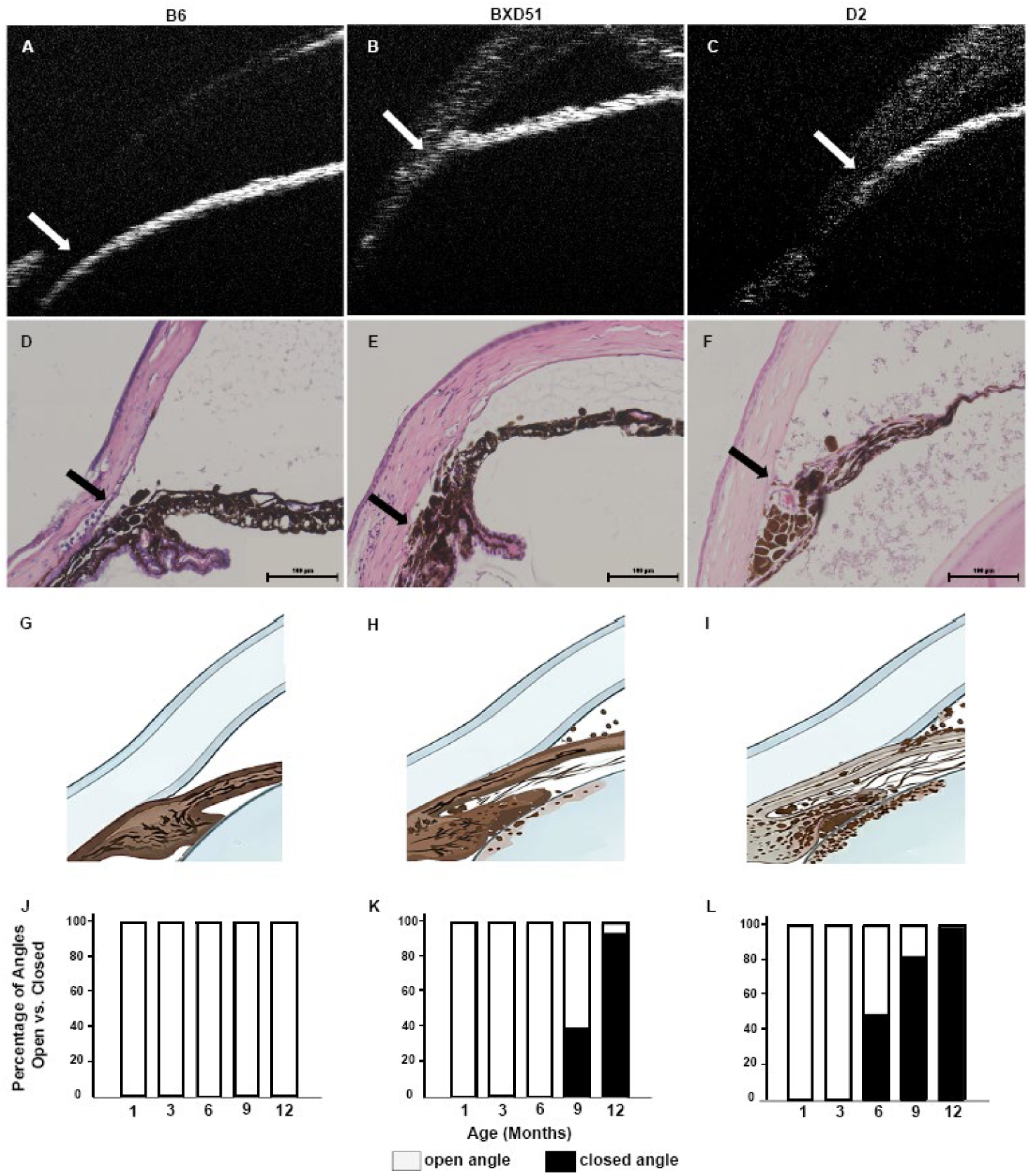
Evaluation of iridocorneal angle morphology and the incidence of angle closure in B6, BXD51, and D2 mice. (A–C) Representative in vivo anterior segment optical coherence tomography (AS-OCT) images of the iridocorneal angle in B6 (A), BXD51 (B), and D2 (C) mice. White arrows highlight the anatomical state of the angle, transitioning from an open configuration in B6 to apposition between the iris and cornea in BXD51 and D2. (D–F) Histopathological sections stained with hematoxylin and eosin (H&E) validating angle status. Black arrows indicate the presence of peripheral anterior synechiae (PAS) and iris adherence in BXD51 (E) and D2 (F) compared with the patent angle in B6 (D). Scale bars = 100 µm. (G–I) Schematic diagrams illustrating iris position and pigmentation loss within the anterior chamber angle. (J– L) Quantitative analysis of the percentage of closed versus open angles across the study duration (1, 3, 6, 9, and 12 months for B6 (J), BXD51 (K), and D2 (L). Data are presented as the percentage of the total number of eyes examined per group. B6 mice maintain a 0% incidence of angle closure throughout the study, whereas BXD51 and D2 mice exhibit age-dependent increases in angle closure, with D2 showing an earlier onset at 6 months of age. Black bars represent the percentage of closed angles, and white bars represent the percentage of open angles.

To track the time course of this endophenotype, we quantified the ratio of open to closed angles at multiple time points (Fig. 3J, K, L). Across the entire 12-month study period, B6 mice maintained 100% open-angle status (Fig. 3J). BXD51 mice followed a delayed course relative to D2; their filtration angles remained open until 6 months of age. From 9 to 12 months, the angles progressively closed, and by the terminal stage the incidence of closed angle due to anterior synechiae reached 95% (Fig. 3K). The D2 cohort showed a faster, earlier onset, with the incidence of angle closure reaching 100% by 12 months (Fig. 3L).

This difference in progression confirms that BXD51 and D2 represent distinct pathological entities, although BXD51 eventually develops the glaucomatous hallmark of angle closure, its more gradual, non-classical course, potentially involving a pigment-dependent component distinct from the classical D2 model, makes it a superior model of studying chronic spontaneous glaucoma.

### BXD51 ON Pathology Mimics Intermediate Human Glaucoma

To examine the molecular changes underlying the observed functional decline, we performed immunofluorescence staining on 12-month-old BXD51 and D2 mice and on human ON tissues, detecting key RGC cytoskeletal and axonal markers: microtubule associated protein 1A (MAP1A), microtubule associated protein 2 (MAP2), and β-III-tubulin (TUBB3) (Fig. 4)[21–23]. Human clinical samples were obtained from the Lions World Vision Institute (Tampa, FL). Demographic and clinical characteristics for these donors are provided in Table 1.

**Figure 4.**
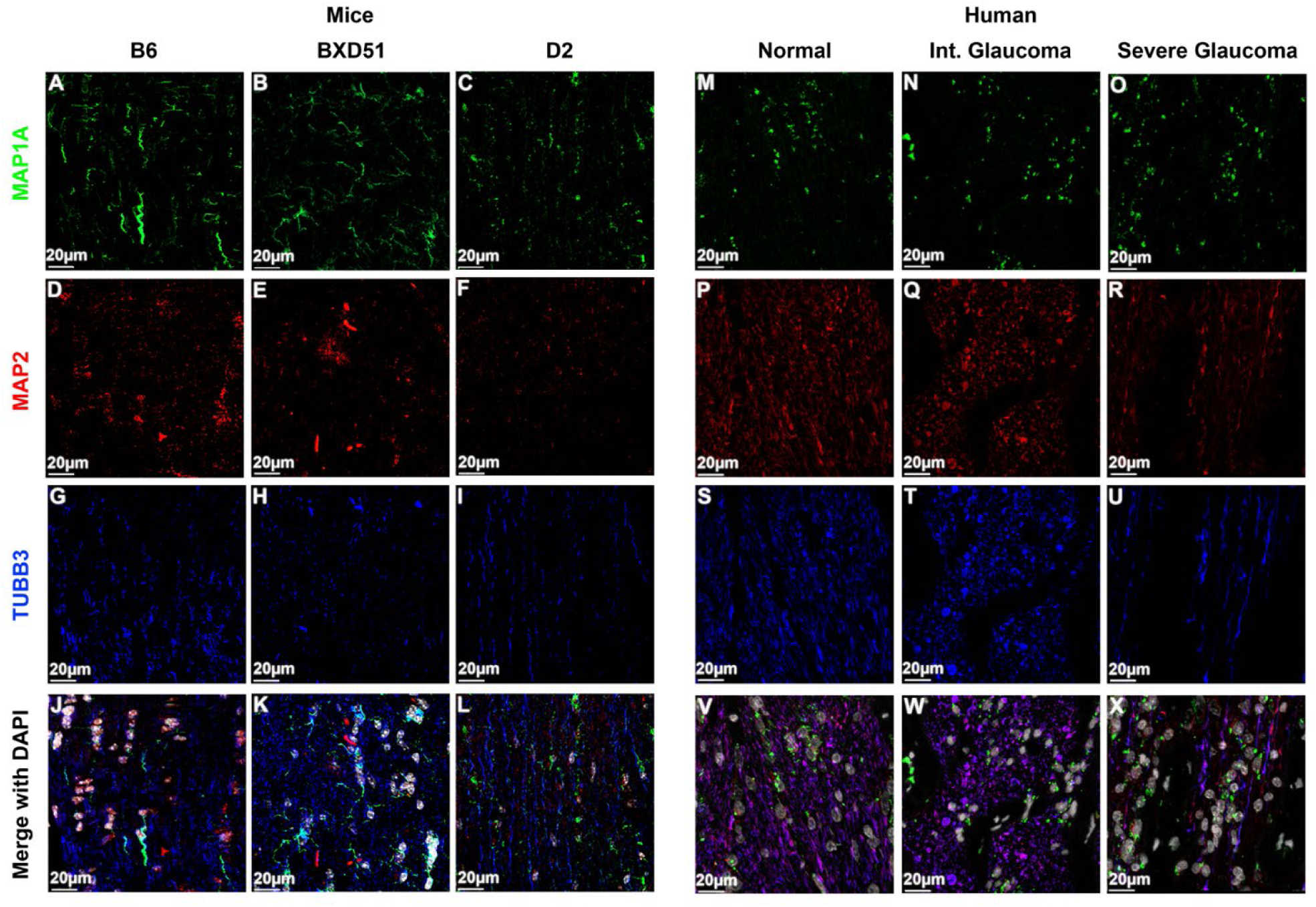
Comparative immunohistochemical analysis of axonal markers in mouse models and human donor optic nerves. (A–L) Representative confocal images of retinal sections from B6 (A, D, G, J), BXD51 (B, E, H, K), and D2 (C, F, I, L) mice. (M–X) Immunostaining of human donor optic nerves from normal controls (M, P, S, V), intermediate glaucoma (N, Q, T, W), and severe glaucoma (O, R, U, X) patients. Sections are labeled with MAP1A (green), MAP2 (red), and TUBB3 (blue), with DAPI (white) used as a nuclear counterstain in the merged panels. Both glaucoma-prone mouse strains (BXD51 and D2) and human glaucoma patients show progressive loss and disorganized distribution of MAP1A, MAP2, and TUBB3 staining compared with their respective healthy controls. Scale bars = 20 µm.

**Table 1:**
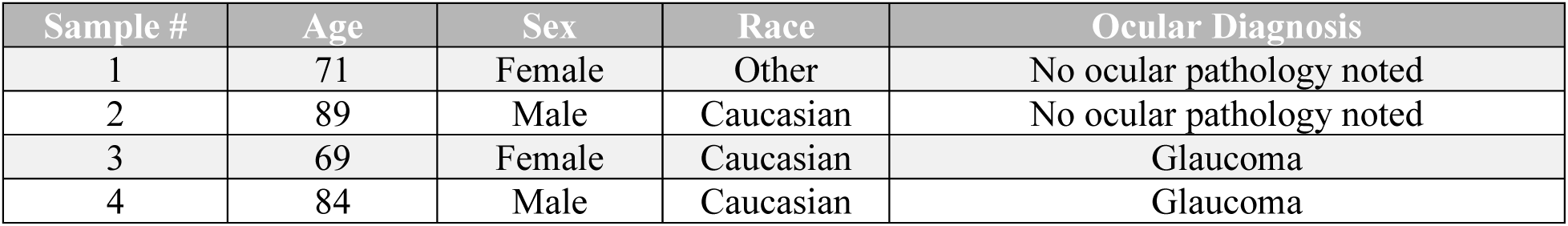
Demographic and clinical characteristics of human tissue donors.

| Sample # | Age | Sex | Race | Ocular Diagnosis |
| --- | --- | --- | --- | --- |
| 1 | 71 | Female | Other | No ocular pathology noted |
| 2 | 89 | Male | Caucasian | No ocular pathology noted |
| 3 | 69 | Female | Caucasian | Glaucoma |
| 4 | 84 | Male | Caucasian | Glaucoma |

In ONs from B6 mice (Figs. 4A, 4D, 4G, 4J) and human donors with no ocular pathology (Figs. 4M, 4P, 4S, 4V), all three markers showed strong, well-organized expression, consistent with a healthy ON population and intact axonal structure. The BXD51 strain showed an intermediate molecular profile (Figs. 4B, 4E, 4H, 4K). Although protein signal intensity in BXD51 retinas was reduced compared with B6, the loss was not as complete as in the D2 strain (Figs. 4C, 4F, 4I, 4L). This preserved but reduced staining pattern in BXD51 mice matches the features seen in human donor eyes with intermediate glaucoma (Figs. 4N, 4Q, 4T, 4W), in which the RGC markers are downregulated and fragmented yet still detectable. In the D2 strain, by contrast, we observed a severe, near-complete loss of MAP1A, MAP2, and TUBB3 signals (Figs. 4C, 4F, 4I, 4L). This widespread depletion parallels the terminal axonal destruction reported in human severe glaucoma (Figs. 4O, 4R, 4U, 4X), where the RGC microtubule network is markedly disrupted.

These histological findings support the value of the BXD51 model from a structural and cytoskeletal standpoint. Whereas the D2 model captures end-stage neurodegeneration, BXD51 provides access to the preceding transitional phases of axonal stress and remodeling. The degradation of the ON cytoskeleton in BXD51 mice was not uniform: TUBB3 showed a comparably severe reduction, but MAP1A expression was relatively better preserved, indicating some resistance during this transitional stress phase.

### Quantitative Analysis of Regional RGC Loss

Prior longitudinal assessments (2015-2021; Supplementary Fig. 1) demonstrated that abnormalities in axonal integrity emerge in the 9 to 13 months interval, with an increasing accumulation of necrotic axons [24], coinciding with the onset of elevated IOP in the BXD51 and D2 strains. To further examine structure of RGCs during this hypertensive phase, we performed a 12-month cross-sectional analysis of the integrity of the RGC complex.

To quantify neurodegeneration in RGC cell bodies, we measured the density of RNA-binding protein with multiple splicing (RPBMS)-positive RGCs across the central and peripheral retinal regions of all three strains (Fig. 5). RGC density showed marked spatial heterogeneity in survival across the retina. Compared with B6 controls (Figs. 5A, 5D), both BXD51 (Figs. 5B, 5E) and D2 (Figs. 5C, 5F) strains showed reduced RGC counts in both central and peripheral regions (Figs. 5G, 5H); however, this loss was not evenly distributed between the two strains. Central retinal loss was comparable between the two genotypes (Fig. 5H), but BXD51 maintained a higher RGC density in the peripheral retina than D2 (Fig. 5G). This pattern contributes to the overall upward, though not statistically significant, shift in whole-retina RGC density in the BXD51 strain (Fig. 5I).

**Figure 5.**
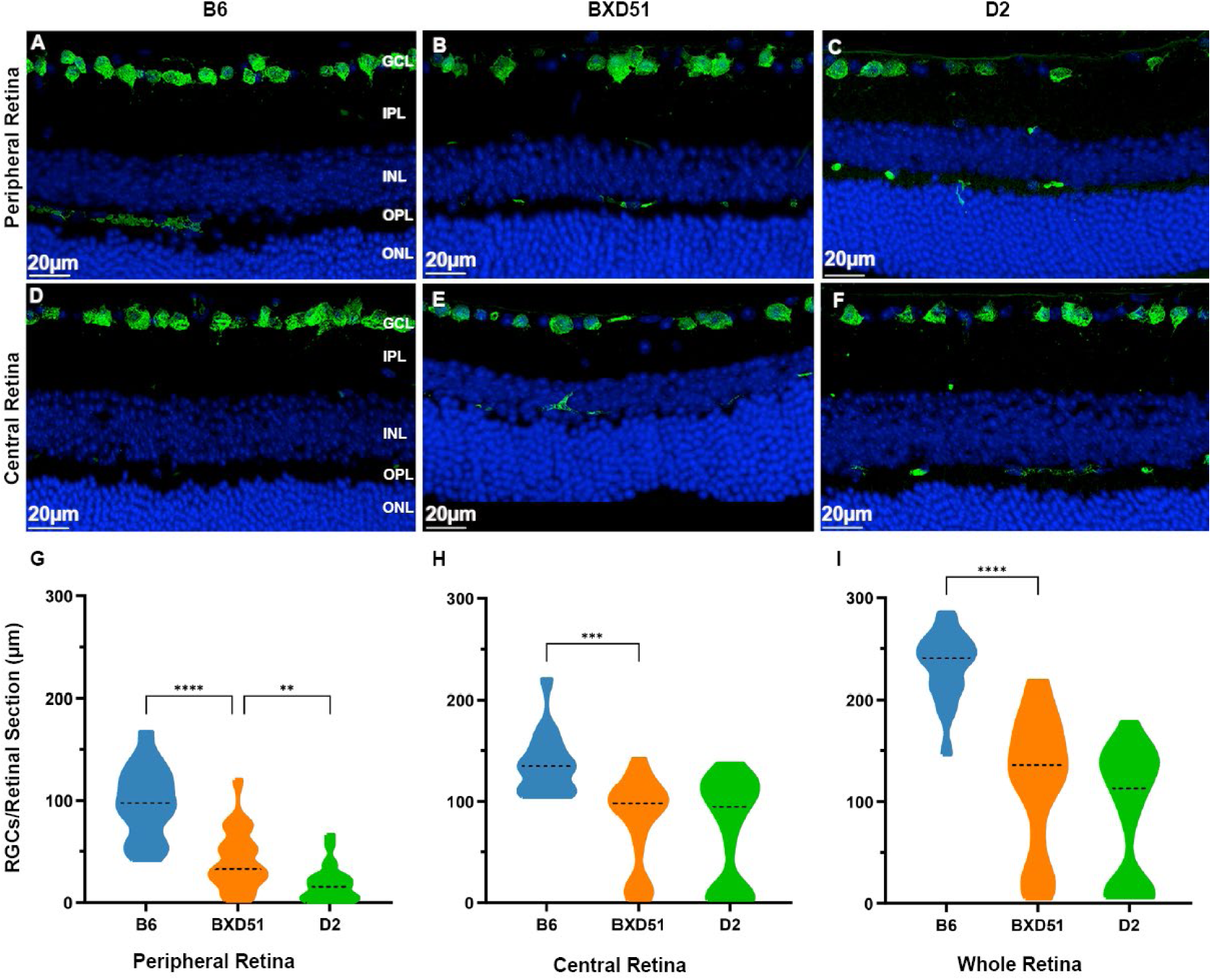
Comparative analysis of retinal cell density and thickness in B6, BXD51, and D2 mice. (A–F) Representative images of retinal sections used for cell density assessment. Micrographs of peripheral (A–C) and central (D–F) retina are shown for the three genotypes. The ganglion cell layer (GCL) is depicted in green, and the inner nuclear layer (INL) and outer nuclear layer (ONL) are visualized via nuclear staining in blue. (G–I) Quantitative analysis of retinal thickness. Violin plots represent the distribution of thickness measurements in the peripheral (G), central (H), and whole retina (I) for B6, BXD51, and D2 mice. Scale bars = 20 µm. \*\**P* < 0.01, \*\*\**P* < 0.001, \*\*\*\**P* < 0.0001.

These findings indicate that central RGC loss is a shared feature of terminal-stage degeneration in both strains, but the spatial distribution of structural preservation differs between the BXD51 and D2 models. This suggests distinct pathological processes underlying neurodegeneration in the two models [25].

### Histological Confirmation of Compartmentalized Degeneration of the RGC complex

To evaluate the structural integrity of the RGC complex—its dendrites, cell body, and axons—we examined the distribution of MAP2 and TUBB3 in vertical retinal sections from B6, BXD51, and D2 mice, as well as from human clinical samples (Fig. 6). In the B6 group (Figs. 6A, 6D, 6G) and normal human donor retinas (Figs. 6J, 6M, 6P), TUBB3 and MAP2 showed intense, continuous labeling within the GCL and the inner plexiform layer (IPL), consistent with a well-preserved RGC complex and dense, arborized dendrites [26, 27]. Consistent with our functional assessments, the BXD51 strain showed an intermediate pathological phenotype (Figs. 6B, 6E, 6H). Although IPL thickness and staining intensity in BXD51 retinas were reduced compared with B6, the retinas retained substantial fragmented MAP2 and

**Figure 6.**
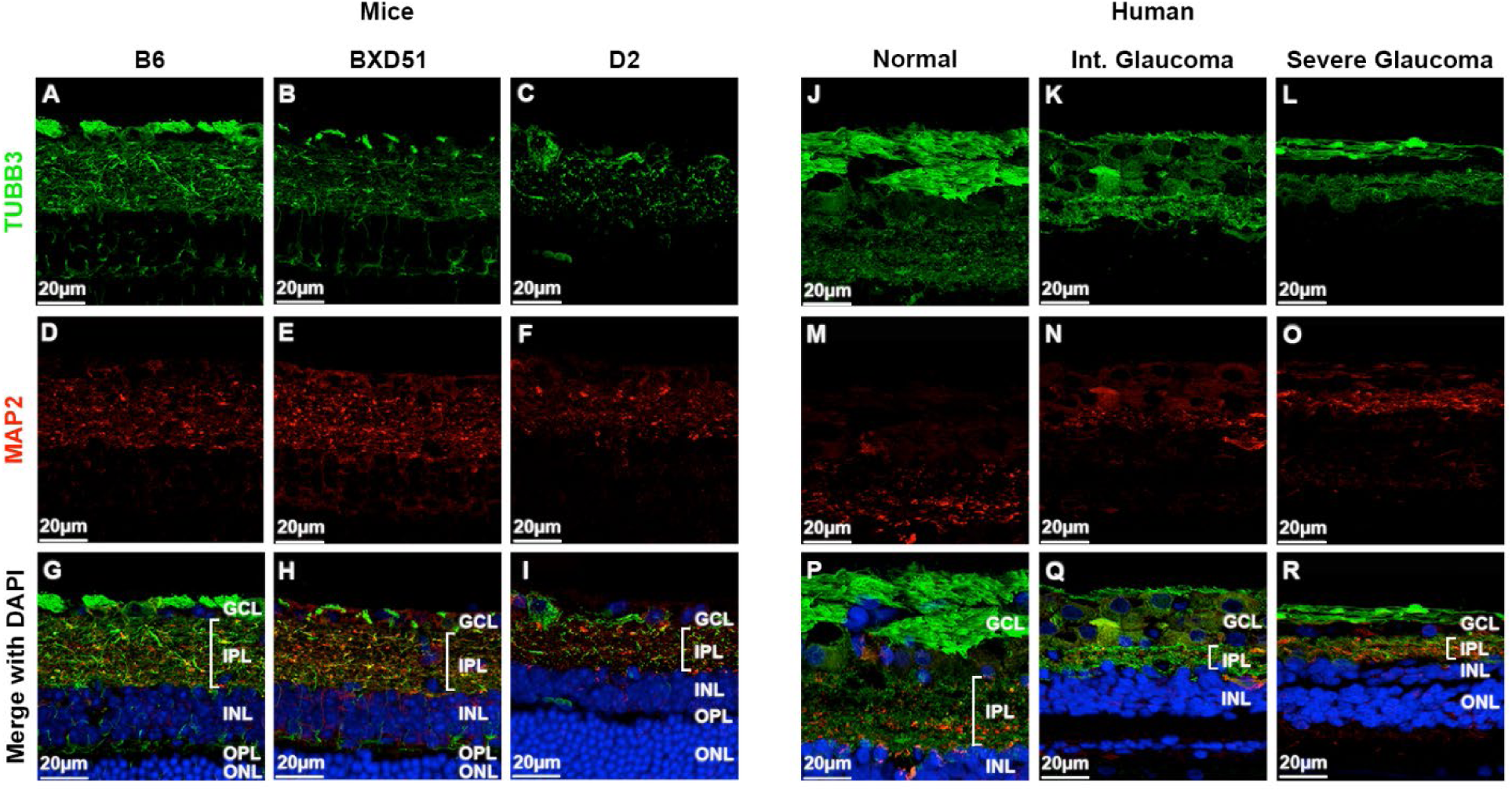
Comparative analysis of retinal laminar structure and marker expression in mice and human donors. (A–I) Immunofluorescence staining of retinal sections from B6 (A, D, G), BXD51 (B, E, H), and D2 (C, F, I) mice. (J–R) Comparative staining of human donor retinas from normal (J, M, P), intermediate glaucoma (K, N, Q), and severe glaucoma (L, O, R) groups. Sections are labeled for TUBB3 (green) and MAP2 (red), with DAPI (blue) as a nuclear counterstain in the merged panels. Key retinal layers are indicated: GCL (ganglion cell layer), IPL (inner plexiform layer), INL (inner nuclear layer), OPL (outer plexiform layer), and ONL (outer nuclear layer). A progressive reduction in the thickness and staining intensity of the GCL and IPL is observed in both glaucoma-prone mice and human glaucoma patients compared with their respective controls. Scale bars = 20 µm.

TUBB3 signal. This pattern, with apparent dendritic remodeling and compensatory hypertrophy under cellular stress, resembles the features of human intermediate-stage glaucoma (Figs. 6K, 6N, 6Q). As in the ON, TUBB3 was substantially depleted in the BXD51 retina. The secondary structural markers, however, showed compartment-specific stability: whereas MAP1A trended toward a milder reduction in the ON, MAP2 was more resistant to degradation within the IPL. The D2 strain, by contrast, showed a pronounced loss of immunoreactivity in both the GCL and IPL (Figs. 6C, 6F, 6I). The thinning of the IPL in D2 mice, together with near-total loss of MAP2-positive dendritic processes, resembles the advanced structural attrition seen in human tissues with severe glaucoma (Figs. 6L, 6O, 6R), in which the synaptic architecture of the inner retina is largely compromised.

Together, these histological findings show a distinct axon-to-soma injury gradient across the visual pathway, consistent with the retrograde neurodegeneration that characterizes human progressive glaucoma.

### Multivariate Discriminant Analysis of Strain Phenotypes

We also analyzed the longitudinal phenotypic and histological data, using five key metrics—IOP, VA, CS, and pERG P50 and N95 amplitudes—in a linear discriminant analysis (LDA) to examine the distinctiveness of and relationships among the three strains at the terminal stage of 12 months (Fig. 7A). The LDA plot showed clear separation among the groups, with the two linear discriminants (LD1 and LD2) accounting for the between-group variance. LD1 captured most of the variance (91.9%) and separated the severely degenerated D2 strain (green squares) from both the B6 and BXD51 groups. The D2 cluster sat far along the positive axis of LD1, at the extreme end of the glaucomatous disease spectrum. The BXD51 strain (orange triangles), by contrast, overlapped only minimally with the B6 control group (blue circles) on the LD1 axis but was much more widely dispersed along LD2. This scatter reflects a delayed progression in which individual BXD51 mice actively resist degeneration.

**Figure 7.**
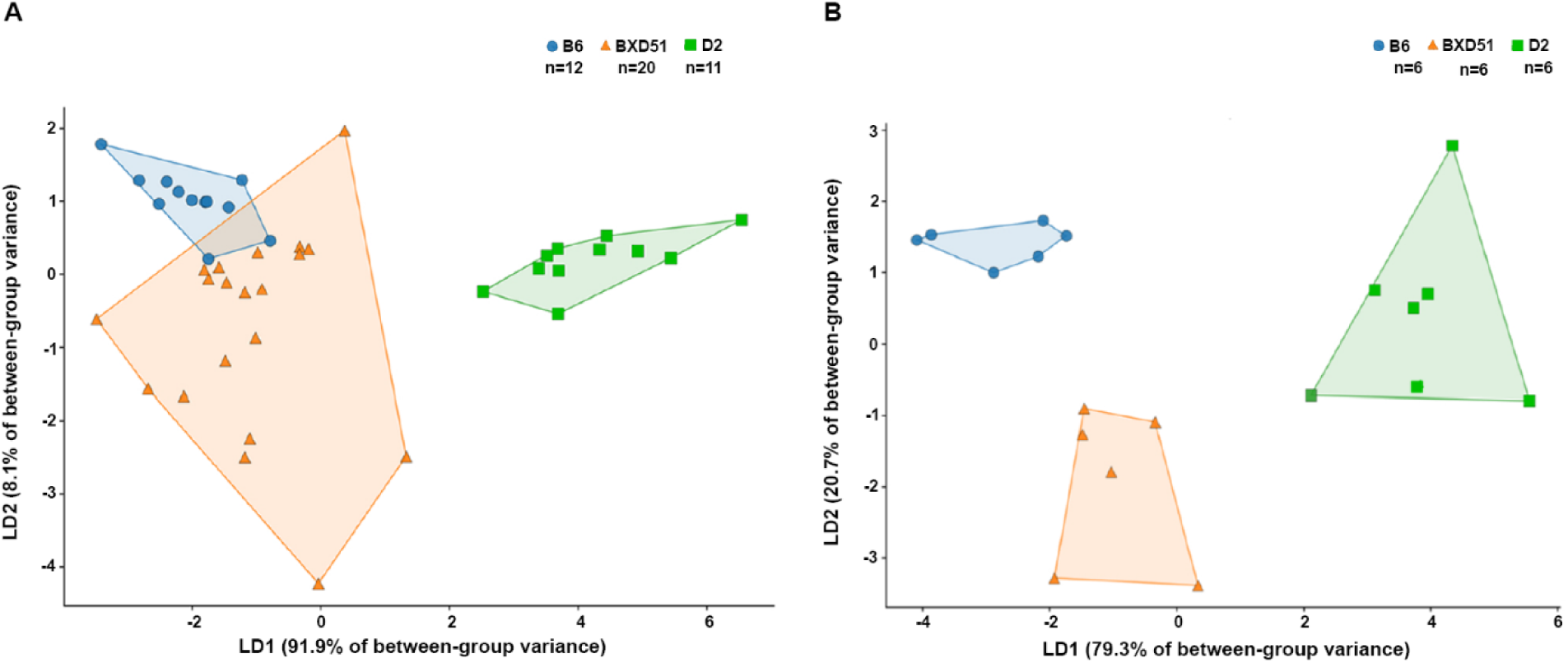
Characterization of the BXD51 clinical and microstructural profiles relative to B6 and D2 mice via linear discriminant analysis (LDA). Clinical functional space (A): Individual B6 (n = 12), BXD51 (n = 20), and D2 (n = 11) mice are projected into the LD1–LD2 discriminant space. LD1 accounts for 96.9% and LD2 for 8.1% of the between-group variance. Included variables comprise standardized measures of IOP, VA, CS, and pERG amplitudes (P50 and N95). The broad, overlapping distribution of BXD51 toward the B6 cluster demonstrates a high degree of shared phenotypic functional resilience during early-to-mid disease stages. Optic nerve microstructural morphometry (B): Individual B6 (n = 5), BXD51 (n = 5), and D2 (n = 5) mice are projected into the LD1–LD2 discriminant space based on optic nerve morphometric variables. LD1 accounts for 79.3% and LD2 for 20.7% of the between-group variance. Included variables comprise standardized measures of mean axon diameter, mean myelin thickness, mean glial area ratio, mean solidity, and mean axon-myelin area. This microstructural space reveals three strictly segregated, non-overlapping genomic clusters. Convex hulls delineate distinct clusters for each genotype, contrasting the shared functional space against the divergent, mechanistically distinct structural signatures underlying health, resilience, and susceptibility.

To determine whether this clinical clustering reflects structural preservation at the cellular level, we quantified the morphometric features of the ON at 12 months. We generated a second, independent LDA model (Fig. 7B) based on five microstructural parameters: mean axon diameter, mean myelin thickness, mean glial area ratio, mean solidity, and mean axon myelin area. With a cross-validated leave-one-out (LOO) accuracy of 100%, this structural model separated BXD51 from both parental strains.

Comparing the two independent LDA models reveals an uncoupling between macro-functional outcomes and micro-structural signatures in these strains. The clinical functional matrix places BXD51 closer to the healthy B6 cohort, reflecting its preserved visual function at 12 months, whereas the morphometric matrix shows a marked spatial shift. This geometric separation indicates that, despite chronic and severe IOP elevation, BXD51 mice do not undergo the structural collapse seen in D2. Instead, they show a localized, distinct tissue-remodeling profile within the ON.

### Co-expression Network Analysis and Core Regulatory Modeling

Given the endogenous neuroprotection suggested by these phenotypic and structural differences, we examined the underlying genomic regulation. Although the BXD51 strain inherits the mutant *Gpnmb* allele from its D2 parent, it retains the wild-type, B6 allele at the *Tyrp1* locus [16, 28]. Working from the hypothesis that this *Tyrp1* B allele acts as a genetic modifier that supports the observed structural and functional resilience, we first extracted the top 500 genes with the strongest expression correlation with *Tyrp1* across the whole-eye transcriptome (from the UTHSC BXD Old Aged Eye RNA-Seq dataset).

Functional enrichment analysis grouped these genes into three main molecular pathways: melanogenesis; PI3K-Akt signaling; and core metabolic networks. To add human relevance, we compared the co-expression network against a panel of human primary open-angle glaucoma (POAG) susceptibility genes and confirmed their expression within our mouse model dataset. From this cross-species alignment, we identified a core group of 64 highly correlated human-derived susceptibility genes. Of these, 8 genes fell within our defined *Tyrp1*-correlated regulatory domain, as shown in the key genes track (Fig. 8A).

**Figure 8.**
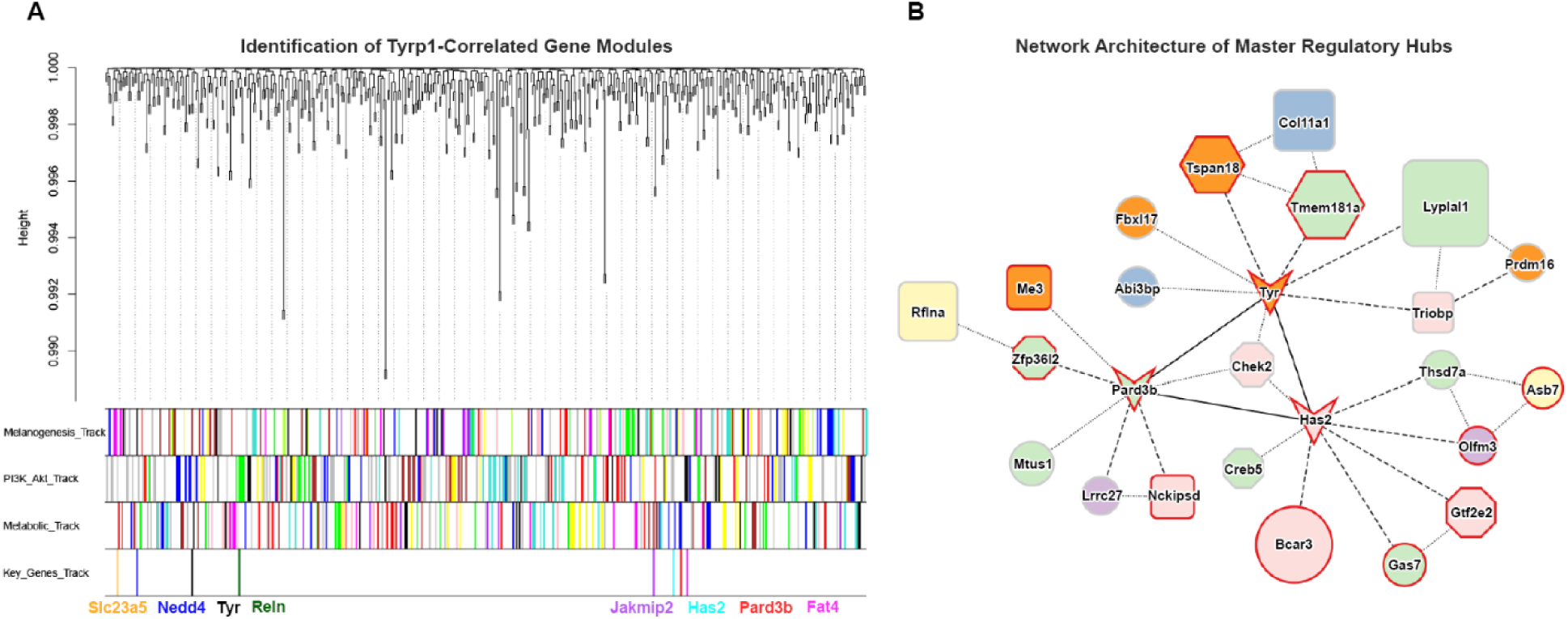
Gene co-expression network driving the unique glaucoma phenotype in BXD51 mice. (A) Hierarchical clustering dendrogram showing gene co-expression modules linked to *Tyrp*1 transcription, delineated into three downstream functional pathways: track 1 (melanogenesis), track 2 (PI3K–Akt signaling), and track 3 (metabolic pathways). Track 4 integrates 64 high-confidence primary open-angle glaucoma (POAG) orthologs prioritized by BXD51 expression significance, highlighting eight overlapping candidate drivers that intersect with the aforementioned functional modules. (B) Network architecture of 22 *cis*-regulatory assist hub genes centered around three master *trans*-hub genes (*Tyr*, *Has2*, and *Pard3b*, represented as V-shaped nodes). Node borders: red outlines denote genes exhibiting distinct transgressive segregation in BXD51 mice. Node size: proportional to the likelihood ratio statistic (LRS) score. Node color (inferred tissue predominance from public databases; see Supplemental Table): orange, retinal pigment epithelium (RPE); blue, homogeneous expression in retina, optic nerve, and RPE; lavender, retina; light green, optic nerve; light yellow, whole eye; pink, trabecular meshwork. Node shape (inferred functional classification from literature consensus): hexagons represent *cis*-triggers; octagons represent regulators; rounded rectangles represent modifiers; ellipses represent satellite genes. Edge type (interaction strength): solid black lines indicate core association axes connecting the three *trans*-hub genes; dashed lines represent proximal associations with high-strength interaction evidence; dotted lines represent peripheral association based on broader statistical co-expression correlations.

To map the regulatory landscape driving the pathobiology of the BXD51 strain, we built a multi-layered interactive network centered on these key regulatory nodes (Fig. 8B). Of the 8 validated candidate genes, 3 nodes—*Tyr*, *Has2*, and *Pard3b* (shown as V-shapes)—demonstrated pronounced transgressive segregation and functioned as central hubs for the network integration of the remaining 22 *cis*-regulated genes. The three master *trans*-hubs and these 22 elements come from the same cohort of 64 high-expression human-homologous POAG susceptibility genes, whose local expression profiles within the BXD51 strain rank at the highest genetic linkage tier under a stringent statistical threshold (peak -log > 4). The network hints a stage-specific flow: a *Tyr*-driven pigmentary insult; followed by *Pard3b*-mediated neuroprotective stabilization; and then *Has2*-dominated structural remodeling. These functional domains are not isolated; they are connected by the regulatory axes linking the core *trans*-hub triangle (*Tyr– Pard3b–Has2*) and share multi-tissue regulators or modifiers.

This *trans*-*cis* coordination organizes the compensatory and pathogenic shifts within the BXD51 strain and provides a molecular explanation for its phenotypic overshoots, such as the mid-stage functional rebound and the late-stage accelerated IOP surge.

## Discussion

In this study, we document the BXD51 mouse strain as a spontaneous, progressive glaucomatous phenotype with anterior segment structural abnormalities, visual functional decline, RGC neurodegeneration, and optic neuropathy. These alterations cover key features of human glaucoma across structural, functional, and cellular domains, supporting the use of BXD51 as a translationally relevant preclinical model[29].

Although BXD51 and D2 strains share the endophenotypes of pigment dispersion and angle narrowing, BXD51 shows a milder, later, and less frequent pathological progression. This suggests that although pigmentary dispersion is a common initiating event for the downstream metabolic and hypertensive cascade, the rate and severity of progression are strain dependent. The angle abnormalities were accompanied by sustained IOP elevation and progressive decline in visual function, including reduced VA and CS. This position is defined not by a simple reduction in severity but by a prolonged compensatory window. The finding that visual deficit precedes overt intraocular hypertension in BXD51 points to an IOP-independent pathogenic trigger. This early stress is often undetectable by iris transillumination at 3 months of age, yet it produces measurable cellular responses and likely stems from pigment-mediated toxicity or iris pigment dispersion-driven cellular stress. The dissociation of VA and CS at 3 months of age may reflect a subtype-specific retinal response and the activation of endogenous protection. Once this buffering capacity is exhausted, visual function deteriorates progressively.

To characterize functional progression in the BXD51 model, we assessed RGC function using the pERG. We identified distinct functional fluctuations, including a biphasic rebound at 6 months of age. Although our historical screenings indicated that significant ON degeneration in this model emerges after 13 months of age (Supplementary Fig. 1), the functional instability we observed at 6 months precedes this window of structural loss. Although these observations come from independent datasets, together they point to a disease trajectory in which early functional decline precedes structural breakdown. This function-before-structure pattern is consistent with established mechanisms in early-stage human glaucoma and positions the BXD51 model as a useful system for studying pre-degenerative checkpoints [30]. Consistent with this, our longitudinal data show a staged functional decline: RGC dysfunction measured by pERG is detectable at 9 months, but the behavioral collapse in VA and CS does not appear until 12 months. This resilience suggests that the BXD51 genetic background engages endogenous neuroprotective mechanisms, such as transient synaptic remodeling or enhanced metabolic compensation, that delay behavioral failure. This extended compensatory phase makes BXD51 a useful model for dissecting the critical disease phases of glaucoma.

At the cellular level, we observed changes in cytoskeletal organization within the ON, reflected in altered microtubule-associated markers. These changes indicate disrupted axonal integrity and may reflect impaired axonal transport, increasingly recognized as a contributor to glaucomatous neurodegeneration[31, 32]. These early axonal and ON-specific changes precede overt pathology in RGC dendrites, consistent with the retrograde degeneration paradigm of glaucoma in which axonopathy precedes soma loss. Following these axonal deficits, quantitative analyses showed significant thinning of the IPL and reduced GCL cell density[33, 34], further supporting the progression of degenerative signals from the ON back to the inner retina. These cytoskeletal changes resemble those seen in human glaucomatous retinal tissue, providing cross-species validation of the underlying pathological processes[35].

Another feature of the BXD51 model is spatially heterogeneous RGC degeneration. In the peripheral retina, we observed a clear genotype-dependent gradient: B6 mice showed preserved structure; BXD51 mice showed intermediate degeneration; and D2 mice presented with the most severe phenotype. In the central retina, by contrast, degeneration converged, and BXD51 and D2 had indistinguishable levels of RGC loss. This suggests region-specific vulnerability, in which peripheral areas retain sensitivity to genetic background while central regions may reach a shared degenerative threshold. Such spatial heterogeneity mirrors patterns reported in human glaucoma and further supports the relevance of this model[36].

A key insight from our multidimensional analysis is the spatiotemporal decoupling between the dynamic clinical functional profile of BXD51 and its underlying micro-structural tissue architecture. Under chronic neurodegenerative stress, clinical function and tissue structure rarely decline along the same linear trajectory. Here, the functional metrics of BXD51—non-linear increased IOP, the fluctuations in VA, CS compensation, and transient pERG overshoots—represent volatile physiological adaptations to ongoing stress. The structural metrics, captured by our axonal and myelin morphometric parameters, change along a more rigid and irreversible path. This uncoupling is visualized by the LDA of the micro-structural matrices. Rather than the tight, uniform collapse seen in the susceptible D2 parent, BXD51 has a broad intra-group dispersion along the primary LD axes. These features indicate that BXD51 does not simply undergo a slower version of the degeneration seen in D2. Instead, its micro-structural matrix aligns more closely with the healthy B6 baseline while occupying a distinct, wide-dispersion cluster. This broad variance suggests that the strain recruits active, heterogeneous tissue remodeling to buffer functional volatility. The LDA framework therefore provides quantitative evidence that the phenotypic resilience of BXD51 is driven by a compensatory structural signature distinct from both parental strains.

A molecular question addressed by our network analysis is why BXD51 maintains overall tissue resilience even though its upstream *Tyr* pigmentary domain contains more transgressive *cis*-elements than the *Pard3b* neuroprotective domain. The BXD51 pathobiology is rooted in the homozygous mutant *D* allele at the *Gpnmb* locus inherited from its D2 parent. In our transcriptomic network, *Gpnmb* acts primarily at a *trans*-regulatory level rather than through a localized *cis*-dominant drop. This indicates that the mutant *Gpnmb* locus does not act as an isolated structural collapse; instead, it functions as a destabilizing genetic background that propagates secondary neuroinflammatory and pigmentary stress across distant functional modules. This heightened sensitivity helps explain why clinical iris pigment dispersion and transgressive molecular activation occur within the *Tyr* anterior domain. This upstream transcriptomic volatility, however, does not translate into early, irreversible tissue destruction. In the classical parental D2 model, severe secondary glaucoma requires a double-hit epistatic interaction between mutant alleles at both the *Gpnmb* and *Tyrp1* loci [37]. In BXD51, this cascade is interrupted because the strain retains the intact, wild-type *B* allele at the *Tyrp1* locus. Acting as a downstream genetic gatekeeper, this protective B allele uncouples the upstream *trans*-acting genetic stress from actual ocular degeneration. By buffering the early pigmentary cascade, this configuration permits an extended phase of sublethal stress and opens a therapeutic window for subsequent multi-tissue compensatory adaptations.

The integrated hub-gene network from our correlation analysis suggests a pathological framework linking the major biological processes identified in BXD51. Functional annotation of the hub genes suggests a sequential organization of these processes. Pigment-associated genes likely act as the initiating trigger of the pathological cascade. These genes connect to inflammatory pathways, which are followed by metabolic stress responses that may delay tissue injury during the early stages of disease. This network mobilization likely represents an energetically demanding, self-limiting defense mechanism. As compensatory mechanisms become insufficient, pathways associated with extracellular matrix remodeling and structural change become more prominent, consistent with the late trabecular meshwork degeneration and rapid IOP elevation observed in BXD51.

Several limitations of this study should be acknowledged. While the integrative bioinformatic approach identifies potential regulatory candidates, the derived correlation matrix reflects functional synchronization rather than direct causal evidence. Further studies, such as single-cell RNA sequencing (scRNA-seq) or targeted CRISPR-Cas9 gene editing, are required to validate these physical interactions. Additionally, although we have established a clear phenotypic and transcriptomic uncoupling, the precise micro-structural chronology bridging the open-angle configuration, initial trabecular meshwork remodeling, and the exact induction point of irreversible neurodegeneration remains to be fully elucidated. Therefore, extended longitudinal studies utilizing real-time, continuous IOP telemetry will be essential to capture the precise temporal sequence of transient fluctuations and to map the non-linear trajectory of functional overshoots against structural decline in greater detail.

In conclusion, the BXD51 mouse strain is a spontaneous, progressive, and spatially heterogeneous model of glaucoma that recapitulates key structural, functional, and cellular features of human disease. The model provides a tractable platform for dissecting disease mechanisms and for evaluating therapeutic interventions across multiple stages of progression. BXD51 serves both as a platform for investigating early molecular mechanisms that link structural stress to cellular response and as a translational tool for developing and testing novel neuroprotective therapies. Understanding these interactions is essential for shifting the clinical paradigm from reactive management to proactive neuro-preservation.

## Methods

### Human Donor Samples

To provide comparative pathological context for the murine phenotypes observed in the B6, BXD51, and D2 strains, post-mortem human donor eyes were utilized in accordance with the tenets of the Declaration of Helsinki. De-identified human eye specimens were obtained from the Lions World Vision Institute (Tampa, FL). The control cohort consisted of two age-matched healthy individuals with no noted ocular pathology, and the glaucoma cohort included two specimens with a confirmed clinical diagnosis of glaucoma. Use of these de-identified specimens was approved or deemed exempt by the Institutional Review Board (IRB) of the University of Tennessee Health Science Center.

### Animals

Three mouse strains (B6, BXD51, and D2) were studied. Foundational breeding pairs for all three strains were originally purchased from The Jackson Laboratory (JAX, Bar Harbor, ME), followed by in-house breeding and aging at the UTHSC animal facility unit. Clinical data (IOP, VA, CS, pERG) were obtained at various ages (1, 3, 6, 9 and 12 months) to capture disease progression. However, histological assessments, including immunohistochemistry (IHC) and ON analysis, were exclusively performed at the 12-month terminal timepoint.

Mice were maintained under conditions of controlled temperature, humidity, and light–dark cycle (12/12 h), with *ad libitum* access to food and water. Data was collected from n≥6 mice, depending on the number of mice available at each timepoint. All procedures including mice were previously approved by the Animal Care and Use review board of the University of Tennessee Health Science Center (UTHSC), Memphis, TN. The protocol also followed the Association of Research in Vision and Ophthalmology (ARVO) Statement for the Use of Animals in Ophthalmic and Vision Research and the guidelines for laboratory animal experiments (Institute of Laboratory Animal Resources, Public Health Service Policy on Humane Care and Use of Laboratory Animals).

### Ocular functions IOP

IOP of the B6, BXD51, and D2 strains were measured using an Icare Tonolab tonometer (Colonial Medical Supply, Franconia, NH) using our previously published methods [18, 38–40]. Data were collected from each strain, both sexes at 1, 3, 6, 9, 12 months of age. IOP measurements were taken between 10 am and 12 pm to reduce the effect of diurnal IOP variation. The mice were anesthetized using a VetEquip rodent anesthesia system (Livermore, CA) with isoflurane (Isoflurane, USP; Covetrus, Dublin, OH). After induction at 2.5%, a maintenance dose of 1.2-1.5% was delivered via nose cone during IOP measurement. Statistical analysis was performed using ANOVA (alpha = 0.05) on Prism 11 software.

### Optokinetic Nystagus (OKN) – VA and CS

VA and CS were obtained from optokinetic nystagmus (OKN) findings. Fully conscious mice were rested on the stage of an OptoDrum OKN machine (Stoelting, Wood Dale, IL). VA data was gathered by fixing the rotation speed on 12°/s and fixing the contrast on 99.72%. CS data were collected by fixing the rotation speed on 12°/s and fixing the cycles on 0.103 cycles/degree with contrast fixed on 99.72%. The statistical analysis was performed using ANOVA (alpha = 0.05) on Prism 11 software.

### OCT

To obtain OCT image, mice were anesthetized using ketamine/xylazine (83 mg/kg ketamine/17 mg/kg xylazine in PBS, pH 7.4) via intraperitoneal injection. Their pupils were dilated with 1% tropicamide. Systane Ultra lubricant eye drops (Alcon, Fort Worth, TX) were applied as needed for corneal lubrication and for clarity. The animals were examined with an Eyemera OCT (IIScience, San Jose, CA) to visualize the iridocorneal angle to assess angle status (open or closed).

### pERG

To access the function of RGCs, mice were dark adapted overnight and anesthetized using the previously stated method. Pupils were dilated with 1% tropicamide. The pERGs were recorded using the Miami pERG system (JÖRVEC, Miami, FL). The P50 wave is a measure of the amplitude from baseline to peak, and the N95 wave is a measurement of the amplitude of the baseline to trough. The data were compiled using Microsoft Excel before it was graphed in Prism 11 software.

### Histological and immunofluorescence analysis

Immunohistochemistry (IHC) was performed post-euthanasia at 12 months of age on samples from B6, BXD51, and the D2 mice of both sexes (n=3-4 mice/strain). All mice were humanely euthanized, and eyes with ONs were removed and placed in a fixative of 4% paraformaldehyde. IHC was performed on retinal and ON sections that were in PBS, pH 7.4 overnight following our published methods [18]. These sections were immunolabeled for markers of RGCs and axons including microtubule-associated protein 1A (MAP1A; MA5-32988, ThermoFisher), microtubule-associated protein 2 (MAP2; MA5-26396, EnCor), and β-III-tubulin (TUBB3; 801202, BioLegend). Secondary antibodies used included goat anti-mouse IgG_1_-AlexaFluor488+ (SA5-10373-AFP488, ThermoFisher), goat anti-chicken IgY-AlexaFluor 555+ (A-21437, ThermoFisher), and goat anti-mouse IgG2a-AlexaFluor647+ (SA5-10370-AFP647, ThermoFisher). Nuclei were labeled with DAPI (D3571, ThermoFisher). Sections were subsequently imaged on a Zeiss 980 laser scanning confocal microscope (LSM) using a 40X objective with 1.3 numerical aperture (NA) or 63X objective with 1.4 NA with or without an associated 0.9X (∼36X) or 1.7X (∼100X) zoom.

### RGC density assessment and retinal thickness measurement

RGCs quantification and retinal laminar thickness measurements were performed on 8 μm-thick paraffin sections labeled for RPBMS (MA5-26396; ThermoFisher), following our established IHC protocol [41]. Secondary antibody used was goat anti-mouse IgG-AlexaFluor488+ (A-32723). Nuclei were labeled with DAPI. Representative sections were visualized using a Zeiss 980 laser scanning confocal microscope.

Data were obtained from B6, BXD51, and D2 mice (n=20-26 eyes per genotype). For each eye, 4–5 non-overlapping retinal sections containing the ON head were selected for analysis to ensure topographic consistency. RGC density and retinal thicknesses were quantified in both the peripheral and central retina, taking into account the central-to-peripheral gradient [42]. During the quantification, measurements from the peripheral and central retinal subsections were recorded separately to analyze regional variations. Statistical analysis and distribution visualization (including peripheral, central, and whole retina data) were performed using Prism 11 software.

### Automated ON morphometrics calculation

Linear discriminant analysis was performed on five standardized clinical measures (IOP, VA, CS, pERG P50, pERG N95) averaged per animal across available ages with complete data for all measures. Bright field images were taken of PPD-stained, resin-embedded ON cross sections from B6, BXD51, and D2 mice (n≥6 nerves per strain). These images were segmented using a deep learning workflow built on AxonDeepSeg: A morphology-based contour extractor merged these axon and myelin masks into a nerve core and applied basic morphology and gradient-plus-spline smoothing to generate whole-nerve contour masks. These contour masks yielded a total cross-sectional area, and glial coverage regions were identified by subtracting axon-myelin masks from whole-nerve contour masks. All segmentation outputs underwent visual quality control review [43–45].

### Cross-Species Bioinformatic Integration and Core Regulatory Network Construction

To elucidate the transcriptional architecture orchestrating the unique glaucoma phenotype in BXD51 mice, a weighted gene co-expression network was constructed using the WGCNA package (v1.74) implement within the R software environment (v4.5.3; R Foundation for statistical Computing) via RStudio [46]. A total of the top 500 genes exhibiting the strongest Pearson correlation coefficients with *Tyrp1* expression profiles were retrieved from the GeneNetwork 2 (https://genenetwork.org) using the UTHSC BXD Old Aged Eye RNA-Seq database and prioritized for downstream module identification. A topological overlap matrix (TOM) was calculated from the adjacency matrix to evaluate network interconnectedness. Hierarchical clustering based on TOM dissimilarity was subsequently performed, and distinct gene modules were partitioned using the dynamic tree-cutting method with the sensitivity parameter set to 2 (deepSplit = 2) and a minimum module size set to 30. To streamline functionally redundant clusters, highly correlated modules were merged utilizing a merge cut height threshold of 0.25 (corresponding to a correlation of 0.75). Functional annotation of the finalized core modules was performed to map cascades, revealing three primary pathological pathways: melanogenesis, PI3K-Akt signaling, and metabolic pathways.

For cross-species translational alignment, human primary open-angle glaucoma (POAG) risk genes were systematically curated from the GWAS Catalog (ebi.ac.uk/gwas) using “POAG” as the primary keyword query. The respective mouse orthologs were subsequently retrieved and intersected with the BXD51 transcriptomic dataset. To filter for strain-specific functional relevance, these candidates were ranked based on their local transcriptomic significance and mapping peak alignment within the BXD51 strain background, isolating 64 high-confidence disease orthologs with a peak -log*P* > 4.

To uncover key drivers bridging clinical relevance and co-expression mechanics, a mathematical intersection was executed between these 64 BXD51-prioritized disease candidates and the top 500 *Tyrp1*-correlated WGCNA module genes, successfully capturing 8 overlapping candidate drivers. The final transcriptional regulatory network consisting of 22 *cis*-regulatory assist hub genes centered around 3 master *trans*-hubs (*Tyrp1*, *Has2*, and *Pard3b*) was visualized using Cytoscape software (v 3.10.4; Cytoscape Consortium). Node topological properties, including likelihood ratio statistic (LRS) scores, were mapped to node sizes (derived from GeneNetwork 2), functional classifications (e.g., *cis*-triggers, regulators, modifiers, and satellite genes) and primary tissue compartments (e.g., retinal pigment epithelium, retina, ON, trabecular meshwork, and whole eye) were conceptually color-coded and shaped based on compiled baseline expression profiles from the EyeIntegration database and established literature consensus (derived from both transcriptomic correlation in GeneNetwork 2 and protein-protein interaction in STRING database), detailed reference data provided in Supplemental Table. Regarding interaction topology, solid black lines indicate the core triangle connecting the master *trans*-hubs, dashed lines represent crosstalk paths bridging distinct functional modules, and dotted lines represent radial links anchoring peripheral satellite nodes to the network core.

### Statistical Analysis

Statistical analyses were performed using GraphPad Prism (version 11) with the significance level set at *α*= 0.05. For the analysis of retinal cell density and retinal thickness, comparisons between genotypes were evaluated using Student’s t-test. For all other quantitative multi-group comparisons including IOP, VA, CS, pERG amplitudes and axonal structural integrity data were analyzed using a one-way analysis of variance (ANOVA) followed by Tukey’s post-hoc test for multiple comparisons. Age-related chronological changes were evaluated using one-way ANOVA or regression analysis, where appropriate. P-values are designated as follows: \**P* < 0.05, \*\**P* < 0.01, \*\*\**P* < 0.001, and \*\*\*\**P* < 0.0001.

## Notes

### Competing Interest Statement

The authors have declared no competing interest.

