## Supplementary figures and images for "BXD51: A Robust and Translational Mouse Model for Studying the Pathophysiology of Glaucoma"

### Supplementary Figure 1

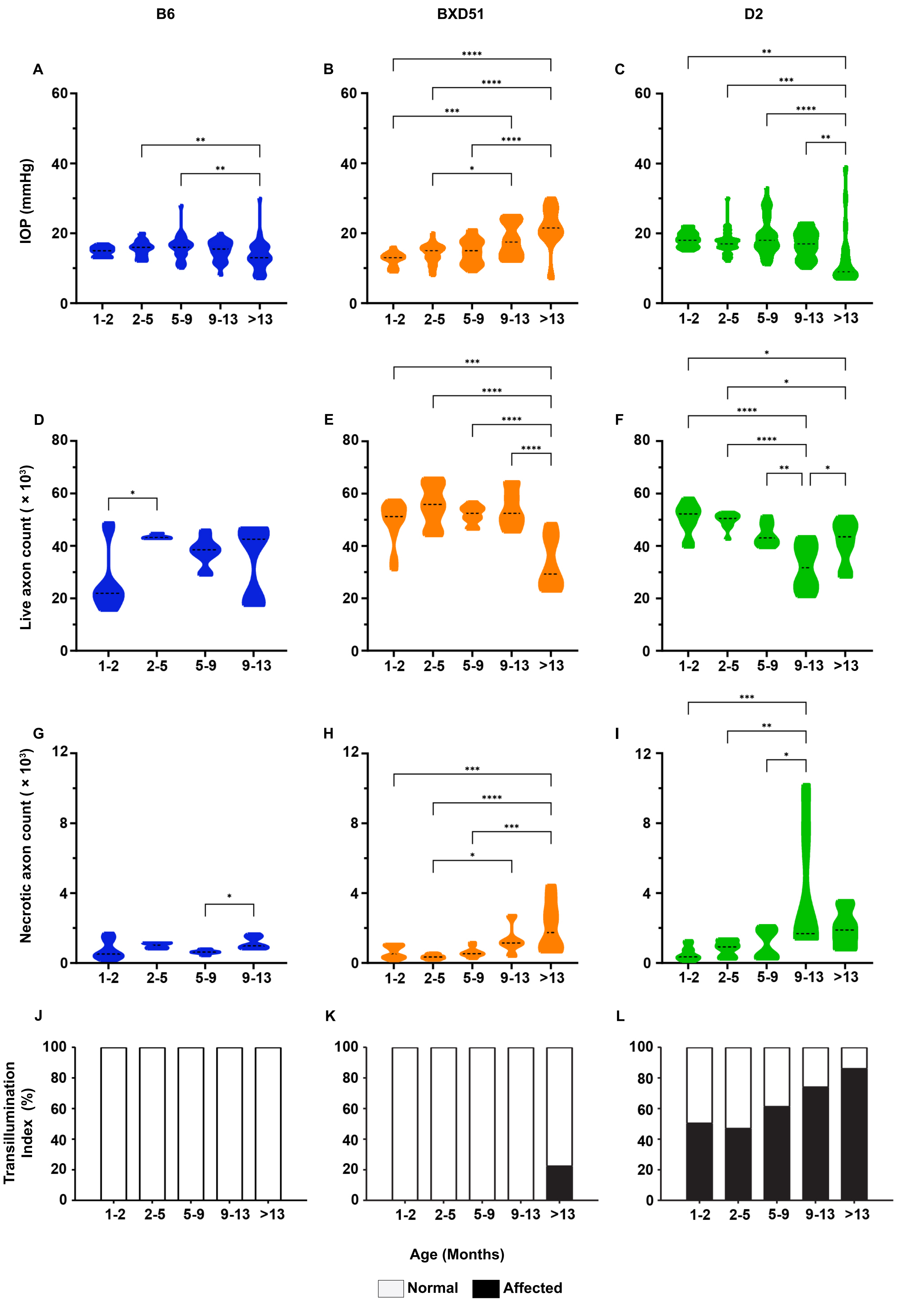
